# How do Cicadas Emerge Together? Thermophysical Aspects of Their Collective Decision-Making

**DOI:** 10.1101/2023.07.31.551222

**Authors:** Raymond E. Goldstein, Robert L. Jack, Adriana I. Pesci

## Abstract

Certain periodical cicadas exhibit life cycles with durations of 13 or 17 years, and it is now generally accepted that such large prime numbers arise evolutionarily to avoid synchrony with predators. Less well explored is the question of *how*, in the face of intrinsic biological and environmental noise, insects within a brood emerge together in large successive swarms from underground during springtime warming. Here we consider the decision-making process of underground cicada nymphs experiencing random but spatially-correlated thermal microclimates like those in nature. Introducing short-range communication between insects leads to a model of consensus building that maps on to the statistical physics of an Ising model with a quenched, spatially correlated random magnetic field and annealed site dilution, which displays the kinds of collective swarms seen in nature.

---

The synchronized spring-time emergence from underground of cicadas of the genus *Magicicada* has been the subject of detailed entomological field studies for over a century [1]. From work documenting the geographic distribution of emergences of 13− or 17− year species [2], to studies of their underground developmental stages [3–5], it is understood that any given brood (group emerging in a particular year) exhibits two types of synchrony; (i) essentially all members emerge precisely in year 13 or 17, and (ii) they do so when the local soil temperature crosses a threshold in that particular year [4].

These observations motivated numerous studies in theoretical population biology to understand the reasons *why* large prime number periods have been selected by evolution, but far fewer studies explaining *how* the two levels of synchrony are achieved. For prime number selection, the hypothesis [6, 7] that limited environmental carrying capacity and predation pressure are responsible was first captured in a mathematical model by Hoppensteadt and Keller [8]. Later models elucidated mechanisms by which single broods occupy disjoint areas [9–11].

These studies do not address how a brood recognizes that it is year 17 (and not, say, 16) and then emerges in a sequence of vast swarms throughout several weeks. The 17 years spent underground by nymphas are divided into 5 *instars*, the duration of which exhibits considerable dispersion (Fig. 1). Despite this spread, cicadas accurately keep track of the passage of years while underground. It is known that after hatching the nymphs burrow below ground and obtain nutrients from the xylem in tree roots [12]. They therefore experience the annual seasonal cycles of the trees, as shown by Karban, et al. [13], who artificially altered the cycles in year 15 to provoke an early emergence, proving that cicadas count cycles and not the passage of time itself. It is unclear how such accurate counting occurs, but it has been suggested [1] that it could involve epigenetic modifications of the kind observed in long-lived plants like bamboo [14]. Similar issues arise in flowin order to flower [15].

**FIG. 1.**
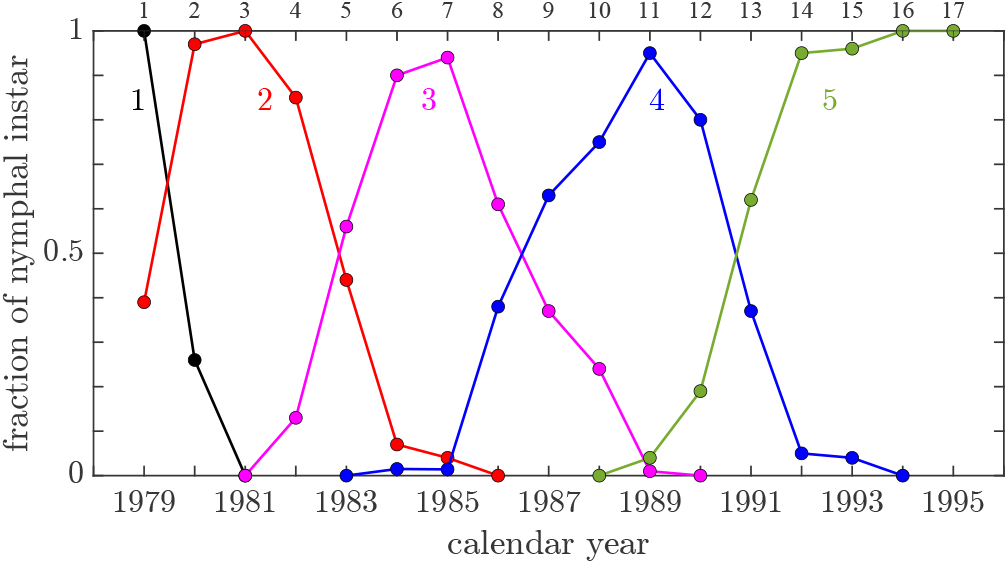
Proportion of cicadas in the 5 instars as a function of time, for one brood. Adapted from Ref. [5].

The issue of swarm emergence in a given year was studied by Heath [4], who found that the day *d*_*c*_ of emergence of 17-year cicadas in any given location is strongly correlated with the local soil temperature reaching the threshold *T*_*c*_ ≃ 18^°^C. This conclusion raises the question of how cicadas can emerge in great swarms in spite of spatially-varying microclimates, their own distribution of body temperatures on emergence [4], and the inherent imprecision of temperature sensing by the cicadas themselves. Here we develop the hypothesis that the thermally-triggered synchronized emergence of cicadas arises in part from short-ranged communication between nearby underground nymphs that allows for collective decision-making. That cicadas are capable of collective behavior by means of communication is evidenced by their acoustically synchronized above-ground choruses [16, 17]. While choruses occur soon after emergence, and it is plausible that the ability to hear underground noise [18] is present earlier, acoustical coupling is but one of several communication mechanisms that may be operating, and our analysis does not depend on the specific means. Collective behavior via communication [19] is found in many contexts: bacterial quorum sensing [20], ant foraging [21–23], and bird flocking [24].

As in other studies of collective behavior [25, 26], our model of decision-making is a random-field Ising model (RFIM) [27, 28], in which quenched randomness arises from microclimates and spins represent the decision. We introduce additional site occupancy variables in order to interpret the simultaneous flipping of many spins (“avalanches”) as swarms. Numerical studies of this model produce swarms like those found in nature.

## Thermobiology of burrowing nymphs

Newly hatched nymphs burrow to a depth *z*_*b*_ ∼ 30 cm that is thought from observations [4] to isolate them from strong diurnal temperature fluctuations. To put this on a quantitative basis, we consider the temperature variations in Ohio, where there is a wealth of data on cicada emergence [4]. Figure 2(a) shows the average daily low and high temperatures at 2 m above ground in Columbus, Ohio [29]. We take these to define a suitable average boundary condition *T* (0, *d*) for the subsurface temperature field *T* (*z, d*) with *z* increasing downward and *d* is time measured in days. These data can be represented by a two-term Fourier series corresponding to a superposition of annual (*a*) and daily (*d*) cycles,

**FIG. 2.**
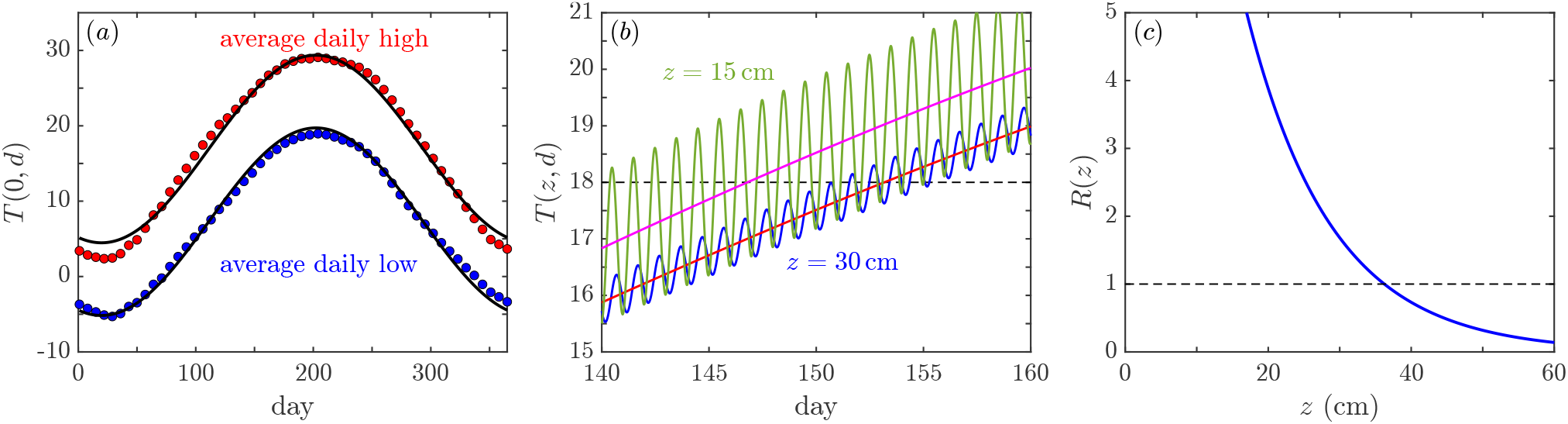
Temperature variations. (a) Daily average low and high surface temperatures in Columbus, Ohio, subsampled weekly, and daily extrema of two-mode approximation (1) (black). (b) Theoretical average subsurface temperature at two depths near *T*_*c*_ = 18^°^C. (c) Noise ratio *R* in (3) versus depth near the crossing day.

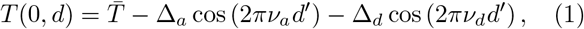

where *d*^′^ = *d* − *d*_0_, with *d*_0_ ≃ 20 (January 20th) being the day of lowest temperatures, with annual frequency *ν*_*a*_ = (1*/*365) day^−1^, daily frequency *ν*_*d*_ = 1 day^−1^, 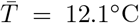, Δ_*a*_ = 12.4^°^C and Δ_*d*_ = 4.8^°^C. We assume the underground temperature *T* (*z, d*) obeys the diffusion equation *∂*_*d*_*T* = *D∂*_*zz*_*T*, for which typical values of the thermal diffusion constant *D* are in the range (0.8 − 10) × 10^−7^ m^2^/s [30]. We adopt the middle of this range *D* ∼ 5 × 10^−7^ m^2^/s= 432 cm^2^/day.

Introducing the scaled time *t* = *ν*_*d*_*d*^′^ and *ϵ* = *ν*_*a*_*/ν*_*d*_, Eq. (1) implies the subsurface temperature field

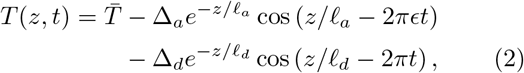

with penetration lengths 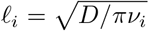 for *i* = *a, d*, with values *ℓ*_*a*_ ∼ 224 cm and *ℓ*_*d*_ ∼12 cm, respectively. Examining the subsurface temperature field at different depths, as in Fig. 2(b), we see that at *z* = 15 cm the within-day oscillations are very large compared to the change in the mean between successive days, whereas at *z* = 30 cm the two are comparable. To quantify the relative size of these two contributions we define *R*(*z*) as the ratio between the root-mean-square daily fluctuations in temperature and the change in the annual trend over one day. Since *ϵ* ≪ 1, we approximate *R*(*z, t*) as

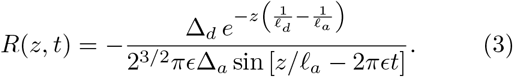

Shown in Fig. 2(c), this ratio decreases with depth, crossing below unity at the burrowing depth *z*_*b*_ ∼ 30 cm. While fluctuations are attenuated relative to the surface, the thermal noise there is comparable to the signal, and thus crossing of the temperature threshold can not be synchronously determined by a population of nymphs, buried at a distribution of depths, acting independently.

### Microclimates and coarse-graining

The above does not account for *lateral* variations in temperature with elevation, tree cover, and solar exposure, which determine the local *microclimate*. As Heath showed, the days of cicada emergences varied with location in a hilly landscape as shown in Fig. 3(a) [4]. Sunny, sparsely forested south-facing slopes have the earliest swarms, with successive swarms typically separated by a gap of several days, disproving the simplistic view that all cicadas in a brood emerge at once within a few days; the entire process within an emergence year may take a month.

**FIG. 3.**
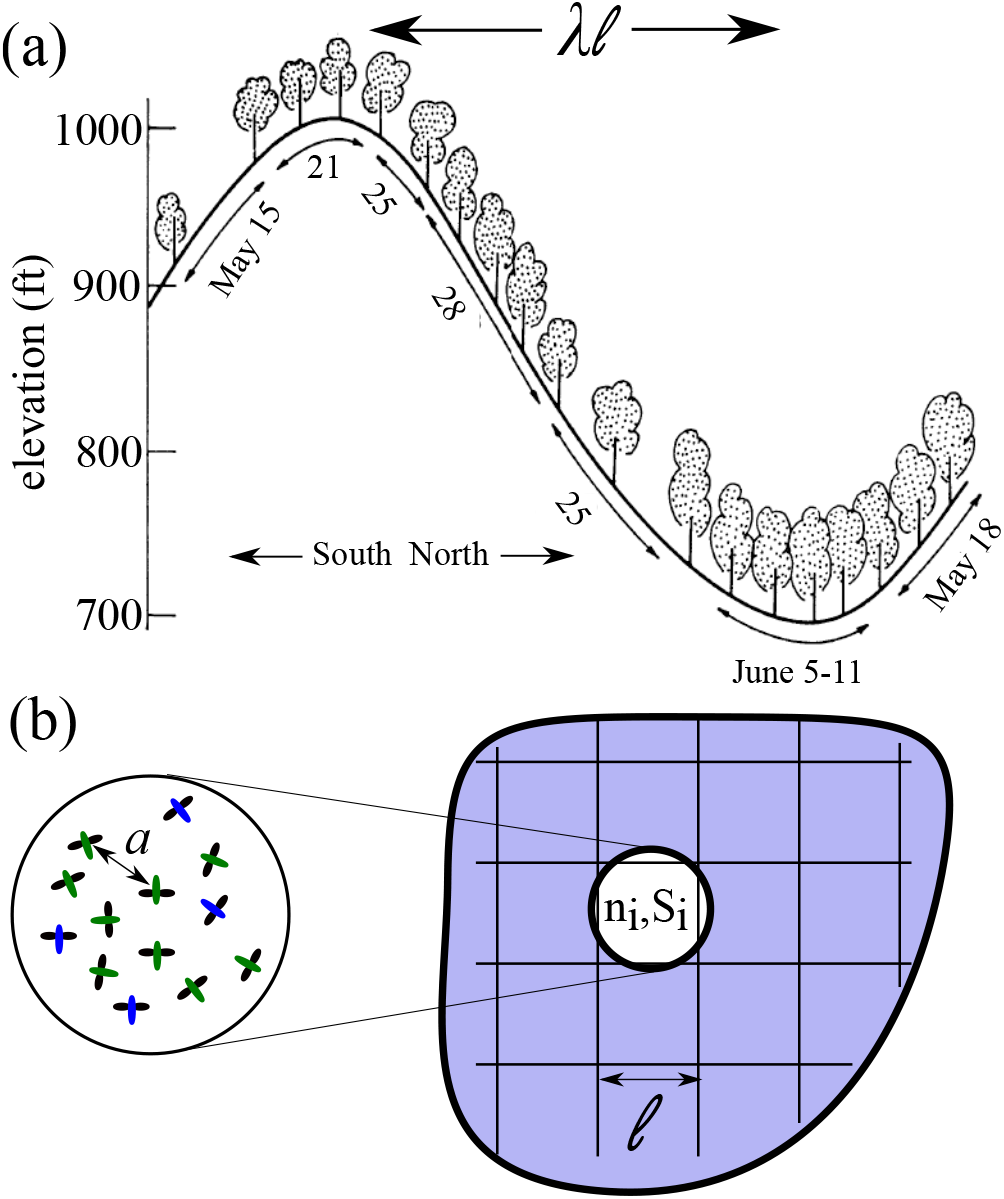
Lateral temperature variations. (a) Topography of an Ohio forest, indicating forest density and dates of cicada emergences. Adapted from [4]. (b) Region of burrowing nymphs, with typical spacing *a*, coarse-grained on the scale *ℓ*. Microclimates are correlated on the scale *λℓ*.

While a full description of microclimate requires accounting for topography, solar exposure, and vegetation, we argue that the net effect of these contributions is that nymphs experience a *quenched, spatially correlated random temperature field*. From our analysis of underground temperatures, we identify the annual penetration length *ℓ*_*a*_ as the smallest scale of that random field which, therefore, serves as a coarse-graining length *ℓ*. The area density *n* of cicadas can reach 10^6^ /acre ∼ 250 /m^2^ [31], with average distance 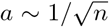 between nymphs as small as 5 − 10 cm ≪ *ℓ* ∼ 2m. We adopt the coarse-grained representation of the population density *n*(**r**) at point **r** in Fig. 3(b), where each subgroup *b*_*i*_ of area *ℓ*^2^ is associated to a site on a square lattice at location **x**_*i*_ = **r**_*i*_/*ℓ* ∈ ℤ^2^ and, as in a lattice-gas description, is assigned an occupation variable *n*_*i*_ denoting if it is empty (0) or occupied (1).

The burrowing depth of nymphs, and the separation of scales *a* ≪ *ℓ* suggest that a natural model of the thermal environment of cicadas involves a two-dimensional temperature field 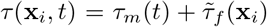, partitioned into a slowly rising mean *τ*_*m*_(*t*) obtained from *T* (*z*_*b*_, *t*) in (2) by averaging over the fast daily oscillations, and a term 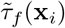 that encodes the fluctuations in the local microclimate. Shifting the origin of temperature to be *T*_*c*_, near the crossing day we may write 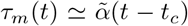, where 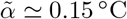. We assume that 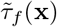 is a Gaussian random field with zero mean and some two-point correlation

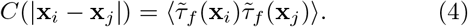

In practice we assume an exponential correlation 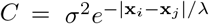 with a single (scaled) length *λ*, where *σ* is the standard deviation of the local field, in the range ∼ 1 − 3 ^°^C. From the topography of Fig. 3(a) and contour maps of the regions studied by Heath, we deduce *λ*∼ 50.

### Model of decision-making

To complete the model by allowing for nymph communication, we introduce a second variable at each site: a spin-like scalar *S*_*i*_(*t*) that characterizes the binary choice at a given time: to remain underground (−1) or to emerge (+1). The decision of group *b*_*i*_ to emerge is determined by the local temperature and the behaviour of other groups in the neighbour-hood *V*_*i*_ (the *q* = 8 nearest- and next-nearest-neighbors of site *i*) via the field ℌ_*i*_(*t*) = 𝒥_*i*_(*t*) + *τ* (**x**_*i*_, *t*)/*σ*, where the temperature has been non-dimensionalized by *σ*, and

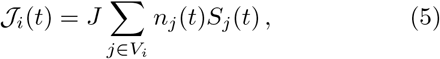

in which we adopt the simplest model with a single coupling *J* throughout the neighborhood. Hence

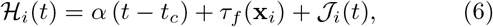

where 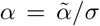, and 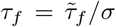 has unit variance. As in previous models of collective decision-making [25, 26], the decision of *S*_*i*_ to flip from −1 to +1 occurs when ℌ_*i*_ becomes positive, as in the “zero-temperature” limit of the RFIM approach. When *J* = 0, each spin flips to +1 when its local temperature field crosses the threshold. When *J* > 0, a subgroup’s decision to emerge is reinforced by occupied neighboring sites that have flipped, a feature that leads to swarms. ℌ_*i*_ plays the same role as the local field in a spin model of magnetization; with the occupation variables *n*_*i*_, the system is random field Ising model (RFIM) with annealed site dilution. In most studies of the RFIM the random field is independent from site to site, but here the microclimates are correlated on scales large compared to the lattice spacing.

The dynamics of decision-making by subgroups is modelled as a discrete-time process in which state variables are updated daily, without resolving the behavior within each day. In numerical studies, we start at *t < t*_*c*_ with full occupancy (*n*_*i*_ = 1, ∀*i*), and with all subgroups choosing to remain underground (*S*_*i*_ = −1, ∀*i*). On each day we iteratively update the spins by the rule 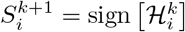 where *k* = 1, 2, … labels iterations, until no more spins flip to +1. We call a *swarm* the set 𝒜 (*t*) of spins that have flipped from −1 to +1 on a given day. The occupancy variables of sites in 𝒜(*t*) are set to zero when the updating rule is complete for that day. The process continues on successive days until the entire lattice is empty.

### Numerical studies

The model (6) has three dimensionless parameters (*α, J, λ*) and the dimensionless system size *L* [32]. Since the tails of *τ*_*f*_ determine the first and last swarms, some 95% of the cicadas emerge over a period of 4/*α* days, during which time the mean temperature sweeps from − 2*σ* to +2*σ* of the random field; setting *α* = 0.3 spreads swarms over the realistic time of ∼ 14 days. Consider first the effect of the coupling *J* at fixed *λ*. Figure 4(a) shows a realization of *τ*_*f*_ (**x**) with *λ* = 30, within which are correlated local “hot-spots” and “cold-spots” in the landscape: like sunny hilltops and shaded valleys. If *J* = 0 (Fig. 4(b)), the swarms are composed of those sites whose random field values fall in intervals of size *α*. As *τ*_*f*_ is Gaussian, the lattice occupancy versus time (Fig. 4(d)) is a discretely-sampled error function 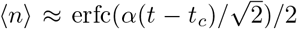. In contrast, when *J* > 0 (Fig. 4(c)), inter-cicada coupling produces large coherent domains. Emptying the lattice involves a smaller number of large swarms, which may be separated by time gaps without activity, as in Heath’s observations [4]. This picture—of quiescent periods punctuated by large emergence events—resembles the avalanches seen in the conventional RFIM, but the event initiation differs due to the daily resetting of the occupancy variables.

**FIG. 4.**
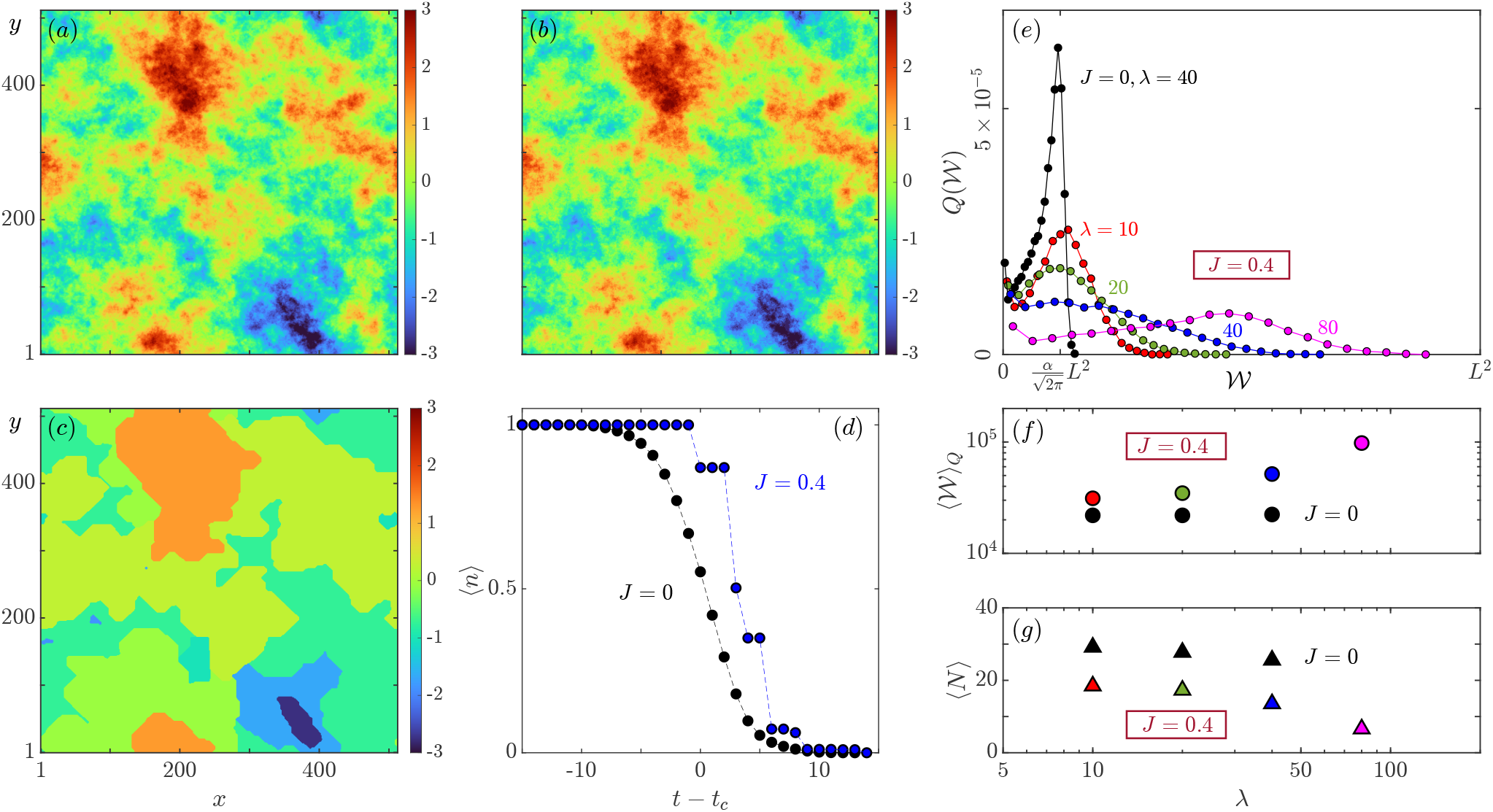
Numerical results with *L* = 512. (a) Realization of the random field *τ*_*f*_ (**x**). (b,c) Composite plots of swarms for *J* = 0 and *J* = 0.4, respectively, color-coded by mean value of *τ*_*f*_ within each swarm, with *α* = 0.3. (d) Occupancy versus time for cases in (b,c). (e-g) Results from averaging over 10^4^ realizations of *τ*_*f*_ for *J* = 0 (black) and *J* = 0.4 (colors). (e) Binned swarm size distribution *Q*(𝒲) for *J* = 0 and for *J* = 0.4 and several values of *λ*. (f,g) Average swarm size experienced by a cicada and average number of swarms versus *λ*. At *J* = 0, the decrease in ⟨*N* ⟩ for *λ* ≿ 40 is a finite-size effect.

Next we examine properties of swarms averaged over 10^4^ realizations of *τ*_*f*_, through the distribution *P* (𝒲) of swarm sizes 𝒲, with mean ⟨𝒲⟩_*P*_ = Σ𝒲*P* (𝒲) and *Q*(𝒲) = 𝒲*P* (𝒲)/⟨𝒲⟩_*P*_, the probability that a given cicada emerges in a swarm of size 𝒲 [33]. We see in Fig. 4(d) that when *J* = 0 the largest swarms occur near *t*_*c*_, where from the form of ⟨*n*⟩ above we deduce the maximum average swarm size to be 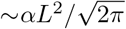. This sharp cutoff is clearly visible in *Q*(𝒲) shown in Fig. 4(e). In contrast, when *J* = 0.4 the pdf *Q*(𝒲) broadens with increasing correlation length of the random field, signifying the existence of ever larger swarms. This is further quantified by examining ⟨𝒲⟩_*Q*_ = Σ 𝒲*Q*(𝒲), the average size of a swarm in which a given cicada emerges. Figs. 4(f,g) show that beyond *λ* ∼ 20, the effect of communication (*J* > 0) is that the average swarm size is larger, and the number of swarms depends more strongly on *λ*. These trends continue for larger *J*.

We have shown that the statistical physics of collective decision-making, quantitatively based on the thermal physics of local microclimates, reproduces key known features of periodical cicada emergences: compact, large swarms spread over several weeks, with temporal gaps between them. Future work could focus on testing the hypothesis of communication between nymphs, and quantifying spatial variations in microclimate and their correlation with emergences. Here we have focused on the synchrony of emergences in year 17. It remains to be seen whether collective decision-making can explain the 13- and 17-year synchrony. Finally we ask: Is there a biological system that exhibits periodic emergences on shorter time scales, allowing for convenient study of this magical phenomenon?

We thank Anne Herrmann for discussions at an early stage of this work and Sumit K. Birwa for assistance with numerical computations.

## Notes

### Competing Interest Statement

The authors have declared no competing interest.

